# Optimizing Primary Human Salivary Stem/Progenitor Cells for Tissue Engineering Applications

**DOI:** 10.64898/2026.05.12.724408

**Authors:** Thaise. C. Geremias, Fabio H. B. da Costa, Nadia G. Mohyuddin, Isabelle M. Lombaert, Mary C. Farach-Carson, Danielle Wu

## Abstract

This work aimed to establish a translationally viable, xeno-free, serum-free platform and protocol for the isolation and expansion of human salivary stem/progenitor cells (hS/PCs) suitable for regulatory qualification and future FDA-approved first-in-human autologous regenerative therapy trials for the treatment of hyposalivation disorders. Parotid gland specimens from non-cancerous regions/tissues were collected from consented surgical patients. Primary hS/PCs were isolated from tissue specimens, cultured in animal-component-free conditions, expanded to produce millions of cells, then enriched for CD44+ stem/progenitor cells by magnetic cell sorting. Normal epithelial purity was assessed using cytokeratins 5/14. Anti-CD133/PROM1 (cancer marker) and anti- fibroblast (clone TE-7) antibodies were used to demonstrate a lack of contaminating cells. Phenotype validation was performed by flow cytometry and immunocytochemistry on both CD44+ sorted and unsorted populations. Senescence-associated beta-galactosidase (SA-β-gal) assays were performed across serial passages (P1–P6). Pluripotency was demonstrated by culture under conditions supporting lineage-specific differentiation. Primary hS/PCs demonstrated consistent expansion and epithelial morphology under serum-free conditions. CD44 expression remained high (>95%) throughout expansion, with negligible detection of CD133 or fibroblast markers, confirming epithelial purity and absence of tumorigenic or stromal contamination. Immunocytochemistry corroborated these expression profiles. SA-β-gal staining revealed only a minor, passage-dependent increase (5–16%) in senescent cells from multiple donors, indicating retention of proliferative potential. Our defined, animal-free culture system supports stable expansion of pure low passage hS/PCs under conditions compatible with good manufacturing practice (GMP).

## Introduction

Cell-based tissue engineering and regenerative medicine applications require the use of well-defined, high-quality implantable cells to ensure safe and efficacious therapeutic outcomes. In addition to identifying a consistent and reliable tissue source of the proper stem/progenitor cells, the cell production process must be carefully controlled whereby all procedures are rigorously defined to deliver consistent cell quality and, ultimately, restore functionality. To meet regulatory standards, cells should exhibit high viability (>90%), robust proliferative capacity during controlled expansion, avoid senescence, and display defined expression of lineage-specific phenotypic markers. Cell populations should be free from microbial or unwanted cell contamination, be non-tumorigenic and not undergo inappropriate differentiation, a problem with the use of induced pluripotent cell (iPC) sources [1]. Primary cells should retain characteristics reflective of their tissue of origin to ensure predictable engraftment and functional integration post-implantation [2].

In the case of the use of human salivary stem/progenitor populations (hS/PCs) for treatment of hyposalivation disorders resulting from radiation therapy for head and neck cancers, we previously developed protocols for the isolation of a therapeutic hS/PC population from surgical specimens of both major and minor glands [3-5] and demonstrated high viability of these cells after implantation into immunocompromised rat and immunosuppressed miniswine models [6, 7]. The cell population we isolate is positive for markers K5, K14 and p63 and derives from the basal cell population of the intercalated ducts of the major glands [8] or a similar population in minor glands [3]. Earlier work showed that hS/PC populations could be differentiated into a secretory, acinar phenotype when hydrogel encapsulated [4].

Across multiple tissues, the hyaluronate receptor CD44 has emerged as a useful surface marker for stem/progenitor cell populations in both murine and human systems [9]. In fact, it was reported that a functional mammary gland could be generated from a single murine CD44+ stem cell [10]. In previous work, we observed that our hS/PC population expresses high levels of CD44 by western blot [11]. Using a different approach, another group showed that human stem cell populations isolated by positive selection using CD34 found that 70% of the isolated cells also were positive for CD44 [12]. The objective of this work was to systematically evaluate the hS/PC populations used in our tissue engineering endeavors, with the goal to evaluate the homogeneity and stability of the cells and to determine if further fractionation of the hS/PC population with CD44 is advantageous.

## Materials and Methods

### Cell isolation and culture

Primary hS/PCs were isolated from normal regions of parotid specimens from consented patients undergoing surgical procedures according to previously described protocol [5]. All procedures performed were compliant with the approved protocols of the Institutional Review Boards at each of the collection sites. Portions of the tissue designated for explant (P0) of hS/PCs were cultured in serum-free complete human salivary medium: William’s E medium (Quality Biological 112014101) supplemented with 10 ng/mL human epidermal growth factor (Gibco PHG0311), 10 µM dexamethasone (Sigma Aldrich D4902-25MG), 1% (v/v) GlutaMax (Gibco A1286001), 1% (v/v) insulin/transferrin/selenium (InVitria 777ITS032), 1 mg/mL (w/v) human serum albumin (Sigma Aldrich A18871), and 1% (v/v) penicillin/streptomycin (Gibco 15140122). Cells were maintained in a humidified incubator at 37°C and 5% CO2; media changes were performed every two days during the primary culture period. When adherent cells achieved approximately 90% confluency, cells were detached with 0.125% (w/v) trypsin 380 mg/L EDTA (Gibco 25200072 + 50% phosphate-buffered saline, PBS) at 37°C and stopped with equal volume of soybean trypsin inhibitor (Millipore Sigma T6522). Cell pellets were separated by centrifugation (233 × g for 2 min at room temperature) and resuspended in complete human salivary media.

### Cell separation using CD44 Microbeads

Cells for each individual sample were subjected independently to magnetic cell sorting using CD44 MicroBeads (anti-human) (Miltenyi Biotec, Germany). For magnetic labeling, adherent hS/PCs were collected as described previously [5] and collected in a cell pellet by centrifugation at 233 × g for 3 min at 4°C. At least 3 × 10^6^ total cells were used for each separation. Pellets were resuspended in 80 µL of adapted magnetic cell sorting (MACS) buffer containing PBS with 2 mM EDTA, 0.5% (w/v) human serum albumin, pH 7.2, and 20 µL of CD44 MicroBeads conjugated with a monoclonal anti-human CD44 antibody (Miltenyi Biotech,130-095-194). The hS/PCs were gently mixed to ensure homogenous labeling and incubated at 4°C for 15 min, protected from light. Following incubation, MACS buffer (2 mL) was added and the mixture was centrifuged at 233 × g for 4 min at 4°C to wash the cells. Supernatants were discarded, and the resulting pellets were resuspended in 500 µL of MACS buffer. Magnetic separation was performed under sterile conditions using an LS Column (Miltenyi Biotec, 130-042-401) placed in the magnetic field of a MidiMACS Separator (Miltenyi Biotec, 130-042-302). The column was pre-rinsed with MACS buffer (2 mL). The labeled cell suspension then was applied to the prepared column. Unlabeled CD44 negative (CD44-) cells passing through the column were collected as flow-through. Each column was washed with sorting buffer (200 µL), and the wash was combined with the flow-through fraction. This combined cell population was collected by centrifugation at 233 × g for 4 min at 4°C, then resuspended in human salivary medium. The bound CD44 positive (CD44+) fraction was eluted by firmly applying the plunger using a 5 mL buffer solution. The CD44^+^ cell suspension then was centrifuged as described above and the pellet was resuspended in human salivary media, followed by incubation at 37 °C in a humidified atmosphere containing 5% CO_2_.

### Flow cytometry

All flow cytometry was conducted at the Flow Cytometry and Cellular Imaging Core at the University of Texas M.D. Anderson Cancer Center (Houston, TX). Each sample was analyzed by flow cytometry independently. Cells were harvested by trypsinization, washed twice with cold PBS and resuspended in adapted MACS buffer. For staining, cells (2 ×10^5^ per sample) were incubated with LIVE/DEAD™ Cell Stain and fluorophore-conjugated antibodies (CD44-FTIC, CD133/2 Antibody-PE and Fibroblast Antibody-APC (see **Supplementary Table 1)** for 30 min at 4°C, protected from light. After staining, cells were washed twice with FACS buffer and resuspended in 500 µL of the same buffer for analysis. Panel Design (BD™ Biosciences) was used to test the compatibility of the fluorochromes by suggesting and blocking those with excessive overlap, according to the selected instrument. Samples were analyzed using a flow cytometer Cytek® Aurora cFluor™5 Laser (Cytek Biosciences, Inc.) and unmixing data was performed by FlowJo® software (Tree Star, version 10). In addition to determining the percentage of cells positive for each marker, the cell size and cell internal complexity (granularity) distribution profiles were analyzed by forward scatter (FSC) vs. side scatter (SSC) plots, respectively.

### Immunostaining for phenotypic biomarkers

Human salivary gland tissues were embedded in optimal cutting temperature (O.C.T) compound (4583, Tissue-Tek) for cryosection (LEICA CM3050). Tissue slices (8 μm in thickness) or cells cultured in 2D monolayers were washed with 1x PBS (without calcium and magnesium), then fixed with 4% (v/v) paraformaldehyde (PFA) for 10 min at room temperature (RT) and washed three times with 1x PBS. Subsequently, samples were permeabilized in 0.3% (v/v) Triton X-100 (Thermo Scientific A16046AP) for 4 × 5 min with agitation and blocked with 10% (v/v) goat serum pH 7.4 diluted in 0.3% (v/v) Triton X-100 for 1 hr at RT with agitation. Samples were incubated with primary antibodies in 10% (v/v) goat serum overnight at 4°C [see **Supplementary Table 1** for antibody descriptions]. Next, samples were washed with 1x PBS (4 × 5 min) and incubated with secondary antibody (Alexa Fluor™ 488 goat anti-rabbit, Alexa Fluor™ 568 goat anti-mouse and Alexa Fluor™ 647 goat anti-mouse) (1:1000, Invitrogen) for staining at RT protected from light. Then, samples were washed four times with 1x PBS and imaged using a Nikon A1R/MP multiphoton confocal microscope and analyzed using NIS Elements software (Nikon Instruments). For staining of cells encapsulated in 3D hydrogels, microstructures first were fixed with 4% (v/v) PFA for 15 min, then subjected to a similar protocol described above except that the time for each step was extended by 50% to guarantee the proper diffusion of fluids through the gels.

### Assessment of senescence

SA-β-galactosidase (SA-β-gal) activity at pH 6.0 was detected cytochemically according to the manufacturer’s instructions (9860, Cell Signaling Technology). Cultured cells were washed once with PBS and fixed in SA-β-gal staining fix solution for 15□min at room temperature. One of the hS/PC populations (M75C P6) was treated with etoposide (ETO) (Cell Signaling, #2200S) using a 10µM dose for 6 hrs and allowed to recover for 72 hrs. Cells then were washed three times with PBS and incubated with SA-β-gal staining solution for 16 hrs at 37□°C. After the overnight incubation, cells were washed with PBS and observed and photographed under a bright field microscope. Total cell numbers and SA-β-gal-positive cell numbers were counted for five randomly selected fields per well. Senescence was calculated as the percentage of SA-β-gal-positive cells per unit area. Results were analyzed by Prism Graph Pad version 10.2.3 software (La Jolla Inc., USA). Statistical analysis was performed by parametric description, obtained data were normal (D’Agostino-Pearson normality test), and compared by ANOVA and Tukey’s post-hoc test (p<0.01).

## Results

### CD44 expression in sorted hS/PC populations

Immunostaining of healthy salivary gland tissue procured from human parotid glands (hPG) confirmed that the CD44 stem cell marker was expressed on cell membranes of acinar structures and stromal interfaces of ductal regions (Fig.1A), whereas K14 was detected in myoepithelial cells encircling acinar cell clusters (red color in Fig 1A). Explants of hPG were used to generate a single-cell suspension of hS/PCs, which became adherent in several T-75 flasks. Colonies grew gradually in size, forming a confluent monolayer of hS/PC (P0) in approximately 10 ± 2 days (Fig. 1B). To investigate the stability of the isolated hS/PCs, cells were enriched for CD44^+^ selection by MACS (Fig. 1C) and cultured over a 2-month period. Homogenous, cobblestone epithelial morphology was seen in all expanded populations across passages (Fig.1D). However, distinct patterns were identified across the post-sort subsets, in which CD44^+^ cells displayed uniform, cobblestone epithelial morphology while CD44^-^ populations presented heterogenous spindle-like and flat cells amongst islands of typical epithelial cells.

**Figure 1.**
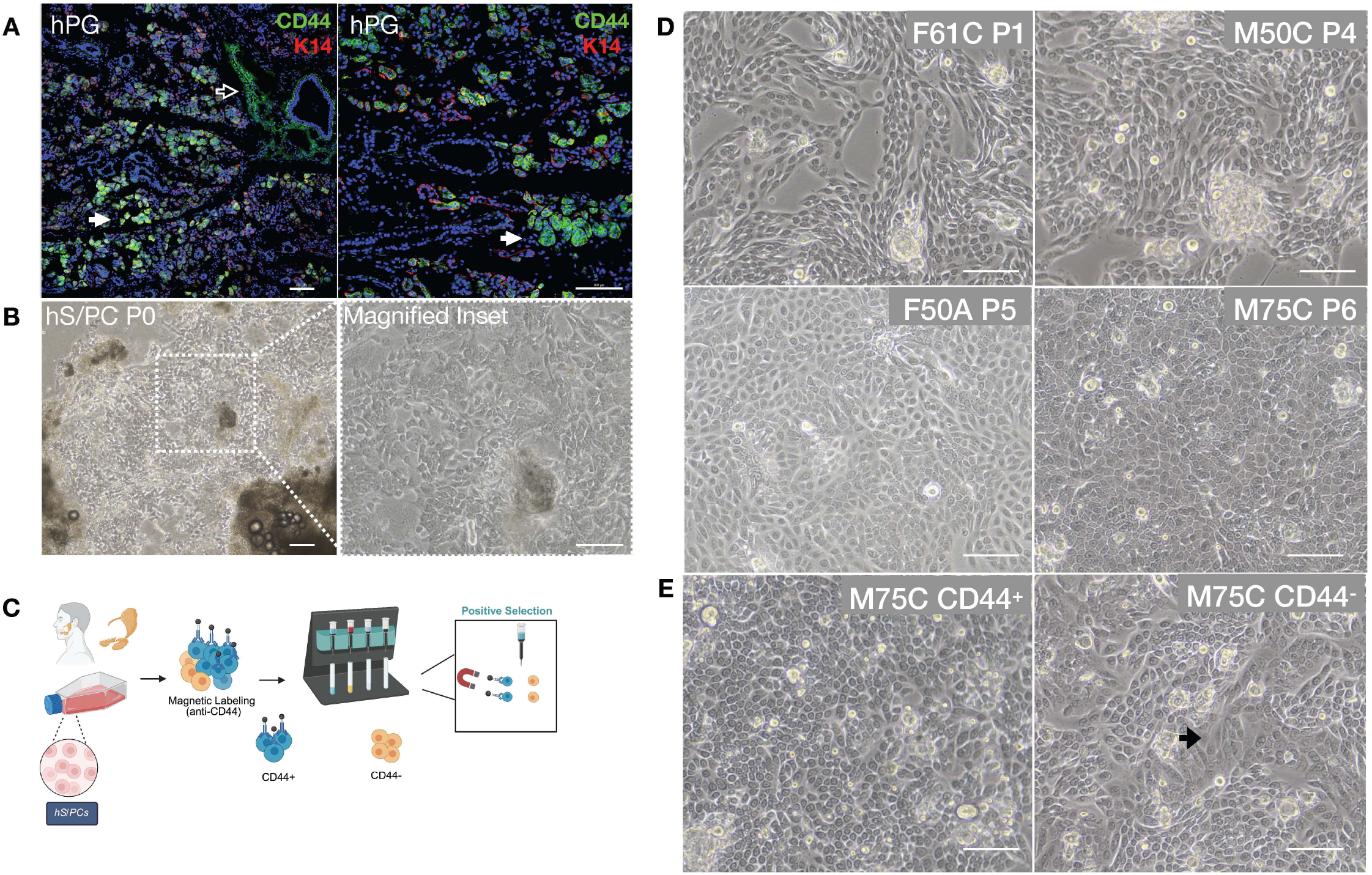
CD44 expression in salivary tissue and primary salivary cell subpopulations. (**A**) Immunofluorescence staining of hPG for CD44 (green) and K14 (red) in tissue from primary tissue specimens; cell nuclei are stained with DAPI (blue). CD44 is expressed on the cell membrane of acinar structures (solid arrow) and stromal regions near the ductal structures (hollow arrow). K14 was localized in myoepithelial cells encircling salivary acini. (**B**) hS/PCs (P0) were harvested from hPG explant outgrowths after enzymatic digestion and re-plated where they formed adherent, small colonies that grew into a confluent monolayer. (**C**) Cell suspensions from monolayers were sorted by magnetic cell-sorting with anti-CD44 microbeads. (**D**) hS/PCs cultured to different passages from four different donors, F61C (P1), M50C (P4), F50A (P5), M75C (P6) showed consistent cobblestone epithelial morphology. (**E**) Sorted hS/PC populations CD44^+^ and CD44^-^. Cultured post-sort for 2 months, the CD44^+^ population displayed a homogeneous cobblestone morphology typical of epithelial cells (left panel). The CD44^-^ cell population (right panel) exhibited cells with a flat, spindle-like morphology (dark arrow) interspersed with typical cobblestone-like colonies. Scale bar: 100 µm in all panels.

### Flow cytometry and phenotype

Flow cytometry was used to evaluate cells derived from multiple donors following explant isolation at passage 0 (P0) and subsequent expansion to confluence at passages P1, P4, P5, and P6, as well as in post-sorted CD44 subpopulations. Analysis of forward and side scatter (SSC) profiles demonstrated that the cultured hS/PCs formed a predominantly homogeneous population, characterized by dense, elliptical distributions (Fig. 2A). This pattern indicates a relatively uniform cellular population, with limited variability in cell size and internal complexity (granularity) among the analyzed samples. Moreover, the absence of widely dispersed or irregular subpopulations suggests high culture consistency and minimal cellular debris or heterogeneous contaminants, supporting the stability and integrity of the isolated hS/PC population under the established serum-free culture conditions. Antibody validation assays performed using colorectal adenocarcinoma Caco-2 cells (P2) for CD133 and human salivary fibroblasts (hSF, P9) for the fibroblast marker as positive controls confirmed specific antibody–antigen recognition under the experimental conditions, thus negatives are true negatives. In Caco-2 cells, the anti-CD133 antibody positively labeled 72.9% of the cell population, confirming adequate sensitivity and specificity for the detection of CD133-expressing cells (Fig. 2B). Likewise, 90.9% of hSF cells were positively recognized by the anti-fibroblast antibody, validating the effectiveness of the fibroblast marker in identifying fibroblast populations. Detection of the respective target antigens in positive control samples demonstrated the specificity of the immunolabeling protocol. Furthermore, these validation data substantiate the consistently diminished fluorescent signal detected across all donor-derived hSPC samples, supporting the conclusion that cancer biomarker CD133 is absent in these cell populations. Phenotypic profiling revealed (Fig. 2C) that the majority of cells expressed high levels of the stem cell marker CD44 (∼99%), with minimal expression of the fibroblast marker (∼1%) and no detectable expression of the cancer-associated marker CD133/PROM1 (0%). These findings confirmed that the expanded hS/PC cultures maintained a largely uniform epithelial phenotype and are effectively devoid of contaminating stromal or malignant cell populations. An increased proportion of fibroblast-like cells was observed at early passage (P1; ∼27%), consistent with residual stromal components originating from the explant isolation process. As expected, fibroblast marker expression was enriched in the CD44^−^ sorted fraction (∼87%), further supporting the specificity of CD44 as a marker for the epithelial stem/progenitor population.

**Figure 2.**
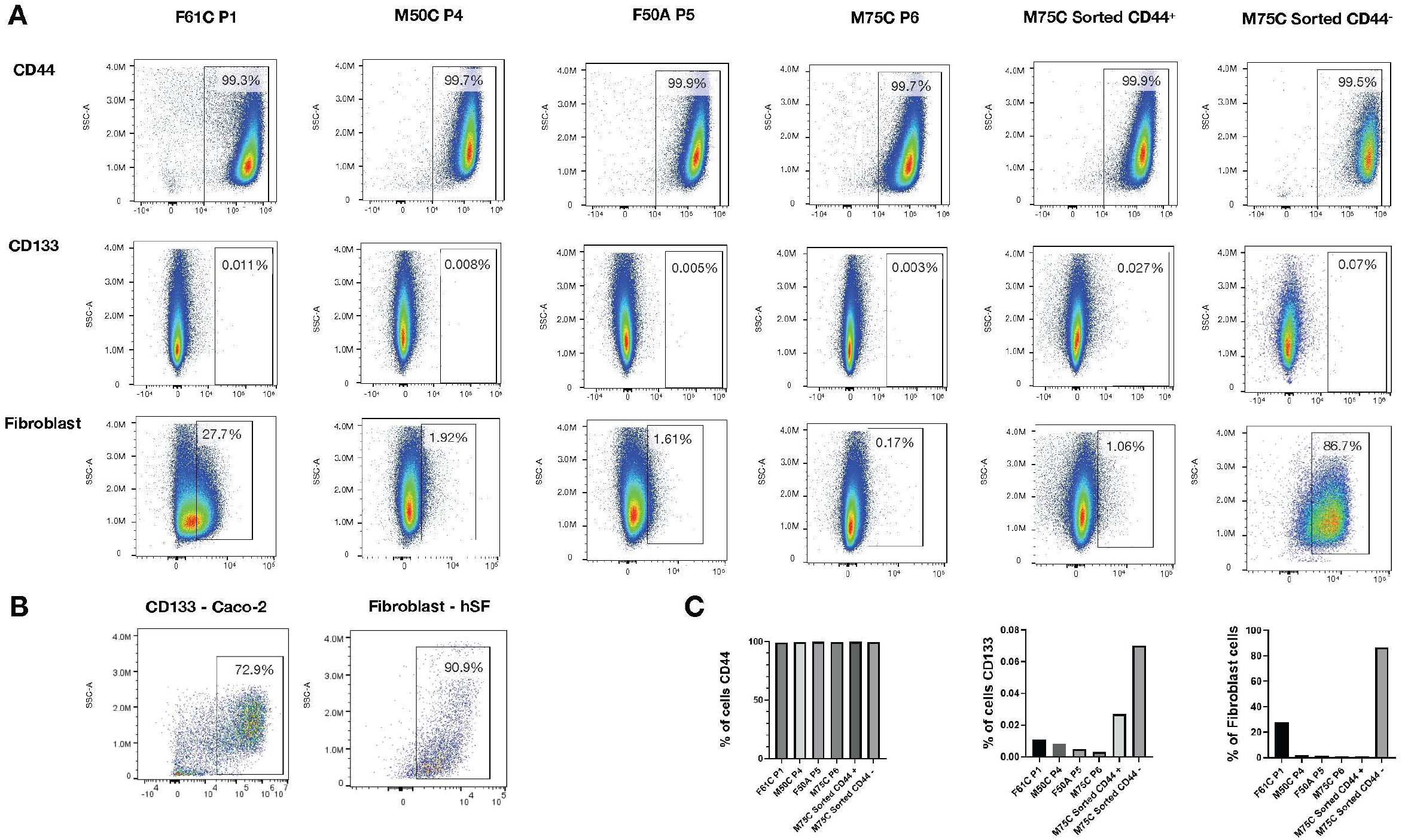
Flow cytometry analysis across different donors. **(A)** F61C (P1), M50C (P4), F50A (P5), M75C (P6) and CD44-sorted populations. Dense elliptical patterns defined all the predominantly homogeneous hS/PCs populations; within the rectangular gates, the presence of high numbers of CD44 expressing cells in all donor specimens was evident; fibroblast antigen expression was only observed in a few cells of early passage then disappeared whereas CD133 was not expressed (P1) even in the more heterogeneous CD44^-^ cultured population. **(B)** Positive controls for CD133, tested in colorectal adenocarcinoma *Caco-2* (P2), and fibroblast marker, screened in primary human salivary fibroblasts (hSF) separately isolated (tested at P9) showed antibody-antigen recognition, thus negative results are true negatives. **(C)** Frequency (%) of CD44, CD133 and fibroblast across different donors.

### Immunostaining and phenotype

Immunostaining results confirmed the expression patterns of CD44, CD133, and fibroblast-associated markers previously identified by flow cytometry across cells derived from different donors (Fig. 3A). CD44 expression was maintained consistently throughout all evaluated passages, indicating preservation of stem/progenitor-associated characteristics during expansion and *in vitro* culture through passage six. In contrast, CD133 expression remained absent across all donor-derived hS/PC cultures, corroborating the flow cytometry findings and suggesting that the isolated cell populations were devoid of neoplastic or tumor-associated cell contamination. In parallel, the fibroblast-associated staining pattern observed in P1 explant-derived cultures and CD44^−^ subset was confirmed, further supporting the phenotypic distinction between the epithelial/stem-progenitor enriched CD44^+^ population and the fibroblast antigen-positive CD44^-^ fraction. Moreover, the sustained maintenance of CD44 expression over extended culture periods further supports the stability of the stem/progenitor phenotype and suggests that the established culture conditions were effective in preserving key cellular characteristics without promoting phenotypic drift or transformation.

**Figure 3.**
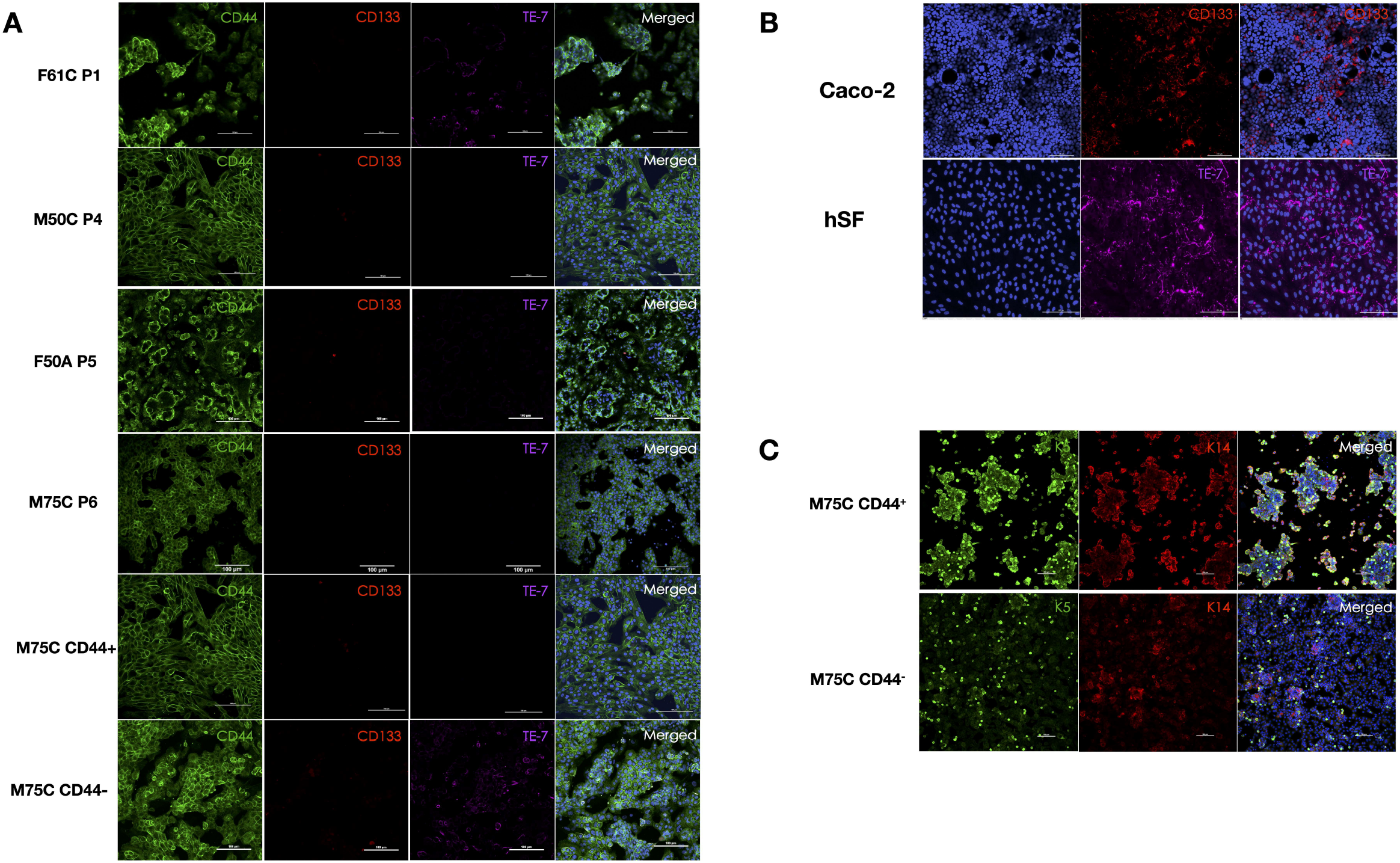
Immunocytochemistry of expanded hS/PC cell populations. **(A)** Sustained CD44 expression (green) was observed across all analyzed passages, demonstrating maintenance of stem/progenitor-related features during prolonged in vitro expansion, including the post-sorted subsets. Conversely, CD133 expression (red) was consistently absent in all donor-derived hS/PC cultures, further supporting the flow cytometry results and indicating that the isolated cell populations were free of neoplastic or tumor-associated cells. Fibroblast-positive cells were identified within early explant-derived cultures (P1) as well as in culture of CD44^−^ cell population, indicating enrichment of fibroblast-like cells within this fraction and highlighting its distinct cellular profile compared with the CD44^+^ epithelial/progenitor subset. **(B)** Immunoreactivity was confirmed using established positive control cells. Caco-2 cells (P2) exhibited characteristic CD133 expression, while hSF (P9) demonstrated positive staining for the fibroblast-associated marker. **(C)** Expression of the epithelial progenitor markers K14 and K5 was detected in both CD44^+^ and CD44^−^ cultured subsets confirming maintenance of epithelial lineage and sustained phenotypic stability during prolonged in vitro culture. Despite expression of cytokeratin in both subsets, K14 and K5 staining intensity was substantially enhanced in the CD44^+^ population, suggesting greater preservation of the salivary stem/progenitor phenotype within this subset. Nuclei are stained in blue (scale bars:100 µm).

To assess staining reliability, a control panel was included (Fig. 3B). As expected, colorectal adenocarcinoma cells (Caco-2) demonstrated positive CD133 expression, whereas human salivary fibroblasts (hSF) showed robust expression of the fibroblast-specific surface marker TE-7. These control results verified proper antibody-antigen recognition and supported the accuracy of the negative and positive staining patterns observed in the experimental hS/PC populations. In addition, expression of cytokeratin 14 (K14) and cytokeratin 5 (K5), well-established epithelial progenitor markers associated with salivary gland stem/progenitor cells, were detected in both CD44^+^ and CD44^−^ cultured subsets (Fig. 3C). Notably, both K14 and K5 immunostaining were markedly more pronounced in the CD44^+^ subset, suggesting that the stronger expression of these epithelial markers in CD44^+^ cells could enhance the preservation of the salivary stem/progenitor phenotype over extended culture periods. Indeed, the sustained expression of these markers in both subsets demonstrates that epithelial lineage identity was preserved and extended in vitro expansion for, at least, two months.

### Proliferative and non-senescence hS/PCs

To assess the senescence status of cultured hS/PCs across expansion, senescence-associated β-galactosidase (SA-β-gal) activity was evaluated at suboptimal pH (6.0) in cells derived from multiple donors at early, intermediate, and late passages. Etoposide (ETO)-induced senescence was included as a positive control to validate assay sensitivity (Fig. 4). At early passage (P1), hS/PCs exhibited minimal senescence, with approximately 5% of cells showing SA-β-gal positivity. These cells displayed characteristic morphological features of senescence, including enlarged and flattened cell morphology, but remained a minor fraction of the population. Intermediate passages (P4–P6) demonstrated a gradual increase in senescence-associated activity, with SA-β-gal positivity ranging from 8.4% to 16.2%, indicating progressive accumulation of senescent cells during in vitro expansion.

**Figure 4.**
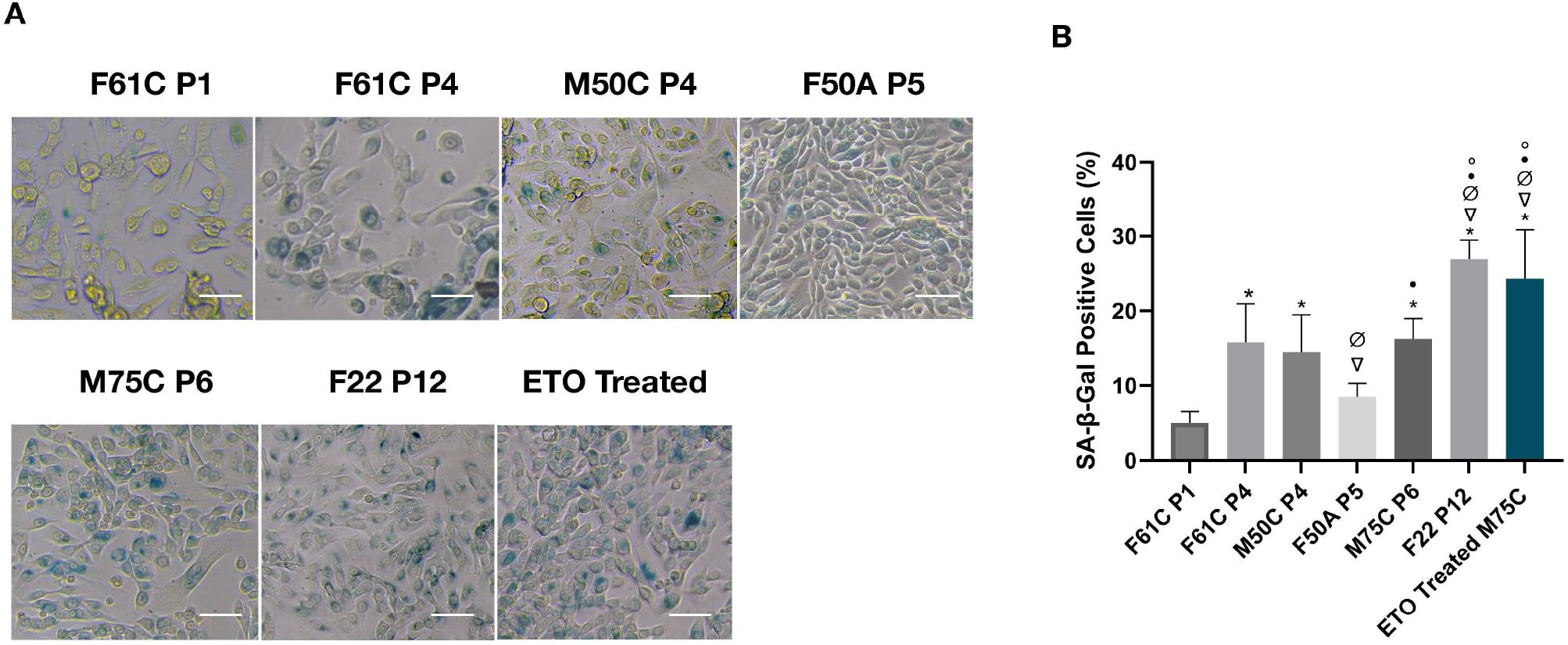
Evaluation of senescence in donor lines across passages. SA-β-Gal staining at pH = 6 was performed in hS/PCs from different donors across passages to assess the degree of senescence. Etoposide-induced senescence served as the positive control. **(A)** Representative images of SA-β-Gal staining (blue color). Scale bar:100 µm. **(B)** The quantification of SA-β-Gal staining is presented as means□±□SD (ANOVA and Tukey’s post-hoc with a significance level of *p* < 0.05). Symbols: (*) represents significantly different when compared to F61C P1; (∇) represents significantly different when compared to F61C P4; (Ø) represents significantly different when compared to M50C P4; (•) represents significantly different from F50A P5; (°) represents significantly different when compared to M75C (P6).

In contrast, late-passage cells (P12) showed a marked increase in senescence, with 26.9% of cells exhibiting elevated lysosomal β-D-galactosidase activity. This significant rise confirms the onset of replicative senescence at advanced culture stages and reflects diminished proliferative capacity and altered cellular function associated with extended passaging. Consistent with expectations, ETO treatment induced a robust senescence response in early and intermediate passage cells, significantly increasing SA-β-gal activity compared to untreated controls. Notably, this effect was less pronounced in late-passage cells, where baseline senescence levels were already elevated, suggesting a ceiling effect in the capacity to further induce senescence in an already aged population.

Collectively, these findings demonstrate a passage-dependent increase in senescence in cultured hS/PCs, with low levels at early stages, progressive accumulation during expansion, and substantial senescence at late passages. This has important implications for defining optimal expansion windows for translational applications, where minimizing senescence is critical for maintaining functional potency.

## Discussion

Distinguishing characteristics of hS/PCs include homogeneous epithelial morphology, high proliferative potential, and expression of biomarkers including p63, K14, and K5 [5, 8]. Surface expression of the stem cell marker CD44 also has been directly associated with human salivary serous acinar cells [13-15], suggesting that enrichment for CD44-positive populations may facilitate isolation of stem/progenitor cell subsets able to maintain and repair salivary gland tissue. In the present study, our serum-free and xenobiotic-free culture system enabled successful isolation and expansion of hS/PCs while supporting manufacturing consistency and safety of the final characterized cellular product.

Flow cytometric analysis demonstrated that cell populations from all donors and across multiple passages exhibited high expression of CD44 (∼99%). Unexpectedly, re-culturing of sorted subpopulations revealed dynamic conversion of CD44 antigen expression, in which MACS-sorted CD44-negative subsets re-expressed CD44 following expansion in vitro. These findings are consistent with published studies describing CD44 as a highly dynamic and microenvironment-responsive molecule whose expression can be regulated by extracellular signals, including growth factors, cell density, and culture media components [16, 17]. These observations suggest that CD44 expression in cultured hS/PCs may reflect phenotypic plasticity rather than a permanently fixed cellular state.

Our culture conditions also effectively enriched epithelial/progenitor populations while progressively eliminating contaminating fibroblasts over serial passages. Initial explant cultures contained approximately 27% fibroblasts; however, continued expansion in our defined medium selectively disadvantaged fibroblast survival and proliferation. By passage 4, fibroblast contamination was reduced to approximately 1%, as confirmed by flow cytometry and immunocytochemistry showing minimal anti-fibroblast marker expression. These findings indicate that the culture system functions as an effective purification strategy for generating a more homogeneous and fibroblast-depleted epithelial population suitable for downstream applications.

Immunophenotypic characterization further demonstrated preservation of epithelial lineage markers throughout culture expansion. Although CD44+ and CD44-sorted subsets displayed slightly different patterns of K14 and K5 expression, both populations retained epithelial morphology and cytokeratin expression during continued culture. Importantly, all analyzed samples demonstrated uniform negativity for CD133. CD133 has been extensively associated with cancer stem cell phenotypes in malignant salivary gland tumors, particularly adenoid cystic carcinoma (ACC), where its expression identifies subpopulations with increased tumorigenic potential, invasiveness, metastatic capacity, and therapeutic resistance [18-20]. The absence of CD133 expression across all donor-derived cultures supports a non-neoplastic biological profile and suggests that the analyzed cells lack molecular characteristics associated with tumor initiation or progression. This observation is particularly important for regenerative medicine applications, where cellular safety and phenotypic stability are critical considerations. Collectively, these data indicate that cultured hS/PC populations maintain stable epithelial and progenitor-associated characteristics during expansion, supporting their suitability for therapeutic and bioengineering applications.

SA-β-Gal staining at pH 6.0 has been extensively used as a marker of cellular senescence both in vitro and in vivo and is known to increase during replicative senescence [21-23]. In the present study, SA-β-gal staining confirmed that late-passage hS/PCs exhibited significantly increased senescence compared with earlier passage groups, although some donor-to-donor variability was observed. Cells derived from donor F50A at P5 displayed lower senescence-associated β-galactosidase activity relative to other experimental groups at similar passages. Notably, hS/PC cultures established from donors of varying ages demonstrated SA-β-gal activity as a function of in vitro replicative age rather than chronological donor age, consistent with previous findings [22].

In addition, low-dose etoposide treatment accelerated the appearance of SA-β-gal staining in hS/PCs, producing senescence levels comparable to those observed in P12 cultures. Similar pro-senescent effects of etoposide have been reported in endothelial cells, where approximately 80% of treated cells exhibited senescence-associated phenotypes [24]. These findings further support the relationship between extended in vitro expansion and progressive acquisition of senescence-associated characteristics in cultured salivary progenitor cells. Future criteria for defining acceptable replicative age for translational applications will require analysis across a larger donor cohort and will likely focus on passages P2–P6 in combination with additional functional and biological activity markers.

## Conclusions

This study establishes a reproducible and rigorously controlled framework for the production and characterization of hS/PCs that is essential for regulatory approval and translational safety. Development of these quality-controlled manufacturing and analytical pipelines represents a critical step toward clinical deployment of autologous stem cell-based salivary gland therapies and supports our ongoing large-animal preclinical studies and planned first-in-human trials.

## Acknowledgments

We acknowledge and thank Dr. Andrew Farach (Houston Methodist Hospital, Houston, TX) and Dr. Quynh-Thu Le (Stanford University) for tissue collection and advice. We acknowledge, as well, all of the research team members from the Farach-Carson and Wu Labs (UTHealth Houston) and Lombaert Lab (University of Michigan).

## Data Availability Statement

The data that support the findings of this study are available from the corresponding author upon request.

