## Supplemental Table 1 for "Optimizing Primary Human Salivary Stem/Progenitor Cells for Tissue Engineering Applications"

***Supplementary Table 1.*** *Antibodies used for Immunocytochemistry/Flow Cytometry*

| *Antibodies* | *Manufacturer* | *Dilution* | *Cat. no.* |
|---|---|---|---|
| CD44-FITC | Miltenyi Inc | 1:400 | 130-113-896 |
| CD133-PE | Miltenyi Inc | 1:400 | 130-113-748 |
| Anti-Cytokeratin 14 (K14) | Abcam | 1:200 | Ab7800 |
| Purified Anti-Cytokeratin 5 (K5) | BioLegend | 1:500 | 905504 |
| DAPI (4′,6-diamidino-2-phenylindole) | Invitrogen | 1:500 | D3571 |
| Fibroblast-APC | Miltenyi Inc | 1:100 | 130-100-133 |
| Anti-fibroblasts Antibody (TE-7) | Novus Biologicals | 1:100 | NBP2-50082 |
| LIVE/DEAD™ Fixable Blue Dead Cell Stain Kit | Invitrogen | 1:40 | L34962A |

Abbreviations: FITC, fluorescein isothiocyanate; PE, phycoerythrin; APC, allophycocyanin.
